# Mammary Epithelial Migration is EMT-Independent

**DOI:** 10.1101/2024.10.17.618667

**Authors:** Jing Chen, Rongze ma, Zhixuan Deng, Yunzhe Lu, Jiecan Zhou, Kun Xia, Ophir D. Klein, Pengfei Lu

## Abstract

Epithelial tissues are traditionally considered immobile, requiring epithelial-to-mesenchymal transition (EMT) for individual or collective cell migration as mesenchymal cells. However, recent studies indicate that the mammary epithelium migrates via mechanisms distinct from mesenchymal collectives. Here, we demonstrate that mammary epithelium does not undergo EMT before migration, as evidenced by cellular and molecular characteristics, including EMT marker expression at the mRNA, protein, localization, and transcriptomic levels. Contrary to established models, cell-cell adhesion is not diminished but is required for epithelial migration. Furthermore, *Snail1* does not repress cell-cell adhesion but instead plays a pivotal role in directing epithelial migration by regulating processes such as the directional movement of individual cells via *Cdc42ep5*. Our findings reveal the existence of both stationary and migratory states within the epithelium, providing new insights into how epithelial tissues transition between these states, with broad implications for organogenesis and cancer biology.

## INTRODUCTION

Epithelium functions as a barrier that covers body surfaces, lines hollow organs and cavities, and forms glands. It is generally considered to be immobile due to its rich expression of cell-cell and cell-matrix adhesion proteins. Classically, it was proposed that epithelium must first transition to a mesenchymal state before migrating as loose individual cells ^1,2^. The process, best known as epithelial-mesenchymal transition (EMT), is thought to be essential for various developmental events, including mesoderm and neural crest formation, and pathologies including fibrosis and cancer metastasis ^3–5^.

Decades of research have since rendered a detailed cellular and molecular description of EMT (Supplementary Fig. 1A). Depending on the context, it is generally believed that various factors, for example TGF-β or ligands of the receptor tyrosine kinase (RTK) family, trigger EMT^6^. This in turn activates a number of transcription factors, especially *Snail*, *Twist*, *Zeb*, etc., which repress expression of epithelial markers including *E*-*Cadherin* (*E*-*Cad*), and occludins, while promoting mesenchymal features, including increased expression of *N-Cadherin* (*N-Cad*), *Vimentin* (*Vim*), and *Pdgfrb* ^5^.

However, inconsistencies with the classical EMT models have also emerged from these studies. For example, E-Cad, whose reduction was considered to be essential for EMT, is instead required for collective migration and cancer metastasis ^7,8^. Moreover, recent computational and fate mapping studies show that only a fraction of EMT markers are activated during certain cancer contexts ^9–11^. This has led to a revised model in which, rather than being in a binary state of either epithelial or mesenchymal, cells exist along a continuum of intermediate states, commonly referred to as partial or hybrid EMT states ^5,12^. Finally, instead of migrating individually as conventionally believed, cancer cells that migrate collectively can disseminate more efficiently and survive better at distinct sites than individual cancer cells ^6,13^.

Recent studies show that mammary gland epithelium migrates in a two-step process that is distinct from all known mesenchymal systems ^14,15^ (Supplementary Fig. 1B). In the first step, stationary mammary organoid undergoes asymmetric cell proliferation and stratification in the front to set up front-rear polarity, i.e., directionality, and generate leader cells, which are essential for powering migration, to become primed organoid ^15^. In the second step, leader and follower cells collaborate to migrate cohesively as a unit. Unlike in mesenchymal systems, where cells are in a fixed position to form a “supracell” ^16,17^, leader cells of mammary organoid epithelium are a dynamic rather than a fixed population in the migration front ^18^. Moreover, they do not send protrusions into the matrix as in the mesenchymal systems^15^. Importantly, the in vitro migrating mammary epithelium resembles at multiple levels the in vivo branching tips, commonly known as terminal end buds (TEBs), including cellular anatomy, architecture, and molecular features ^15^. Indeed, genetic ablation of the tissue polarity gene *Lgl1* not only blocks epithelial migration in vitro but reduces branching in vivo as well ^19^, suggesting that collective migration occurs in the TEBs as an essential part of normal epithelial branching.

This previous work showing that mammary epithelium migrates distinctively from mesenchymal cells has led us to hypothesize that it does not transition to a mesenchymal state before migration onset. Here, we test this hypothesis by systematically examining the cellular and molecular events essential for mammary epithelial migration.

## RESULTS

### Mammary Epithelium Does Not Show Typical EMT Signatures During Collective Migration

First, we examined whether TGF-β, one of the best studied EMT promoter ^5,20^, facilitates epithelial migration. To this end, we added TGF-β at 0 ng/ml or 5 ng/ml concentration to the medium and subjected mammary organoids to the migration assay. Surprisingly, the addition of TGF-β not only did not promote migration, e.g. by shortening the priming stage where organoid was not migratory or speeding up subsequent migration, it completely blocked it (Figure 1A-B).

**Figure 1:**
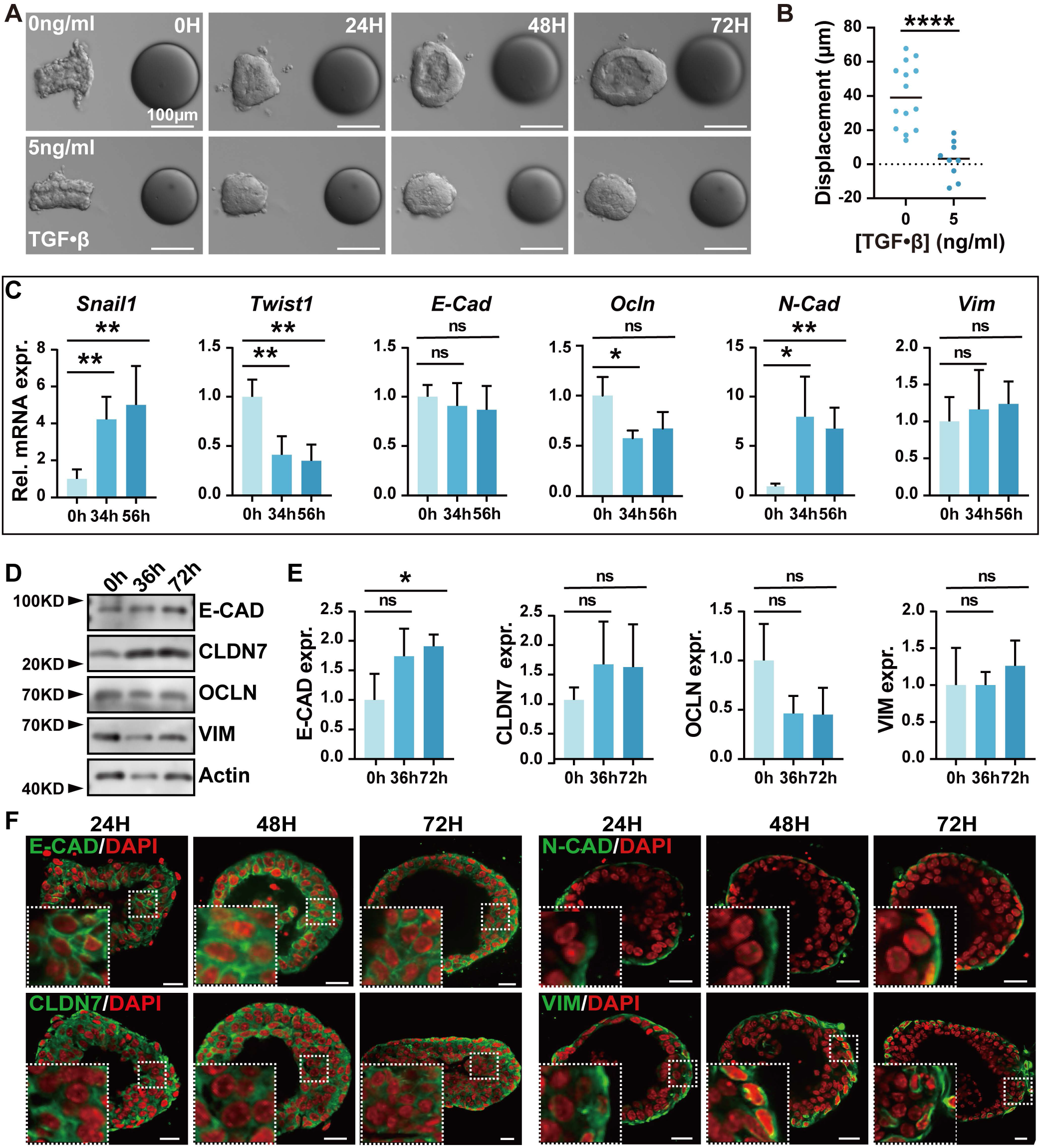
Mammary epithelium does not show typical EMT signatures during collective migration. (**A, B**) Effect of EMT promoter TGF-β protein on FGF10-induced epithelial migration. TGF-β was added to the medium at the concentration indicated (**A**). Effect of TGF-β at 0 ng/ml (n=14) and at 5 ng/ml (n=10) was quantified (**B**). (**C**) Relative mRNA expression of a panel of EMT markers as detected by qPCR at different stages of mammary epithelial migration. Note in the 3D in vitro migration model, mammary epithelium is apical-basal (AB) polarized and stationary at the 0-hour stage, has transformed AB- polarity to front-rear (FR) polarity in the front by the ∼32-36-hour stages, after which it actively migrates. (**D, E**) Protein expression as detected by Western blotting (**D**) and quantification (**E**) of select EMT markers at different stages of epithelial migration. Note β-actin was used as an internal control. (**F**) Protein expression and localization of selected EMT markers as detected by Immunofluorescent staining at different stages of epithelial migration. Insets show a high magnification of the area with a white-dotted frame. N=3. Scale bars: 20 µm. Data are mean ± SD. Statistical analysis was performed using unpaired Student’s t test. *p < 0.05; **p < 0.01; ****p < 0.0001; n.s., not significant. Abbreviations: Rel, relative; expr, expression.

Next, we assessed how EMT markers change their expression during the epithelial migration process. Using qPCR reactions, we determined the mRNA expression of a typical panel of EMT markers, including the transcription factors *Snail1*, *Twist1*, and *Zeb1*, cell junction genes including *Cdh1* (encoding E-CAD), *Cdh2* (encoding N-CAD), *Occludin* (*Ocln*), *Claudin7* (*Clnd7*), the cytokeratin gene *Vim*, and *Pdgfrb* at the start of the experiment (0 hour), migration onset (34 hour), and during active migration (56 hour). Interestingly, except for *Snail1*, *Ocln*, and *N-Cad*, most of the other marker genes did not show an expected change in mRNA expression; in fact, *Twist1*, and *Pdgfrb* even showed an opposite change, i.e. decreased mRNA expression (Figure 1C and Supplementary Fig. 1C).

Using Western blotting, we examined the protein expression of some of EMT panel members. We found that, largely consistent with the above qPCR expression data, protein expression of E-CAD went up, but CLDN7, OCLN, or VIM did not show a significant change of protein expression during epithelial migration (Figure 1D, E). Next, we used immunofluorescent confocal microscopy to examine protein expression of the epithelial markers E-CAD and CLDN7, and the mesenchymal markers N-CAD and VIM during migration (Figure 1F). We did not observe an obvious amount change of protein expression of either the epithelial or mesenchymal markers. Interestingly, we found that both N-CAD and VIM were enriched in the basal epithelium and excluded from luminal cells throughout the migration stages (Figure 1F).

To confirm whether these presumed mesenchymal markers are preferentially expressed in these two subtypes of mammary gland epithelium, we performed qPCR reactions using cDNA templates from basal and luminal cells from the 7-week branching mammary epithelium. We confirmed that *N-Cad* and *Vim* were enriched in basal cells with little expression in luminal cells (Supplementary Fig. 1D). Likewise, other EMT markers including *Twist1*, *Zeb1* and *Pdgfrb* were also enriched in basal cells, whereas *E-Cad*, *Ocln*, and *Cldn4* were enriched in luminal cells (Supplementary Fig. 1D).

Together, these data show that mammary epithelium does not show typical EMT signatures during collective migration. Moreover, the observation that EMT markers are already differentially expressed in the subtypes of stationary mammary epithelium and that this expression pattern persists throughout different stages of mammary epithelial migration calls into question their accuracy as EMT markers.

### Mammary Epithelium Does Not Undergo EMT at the Transcriptomic Level

To gain a comprehensive view of the molecular events during epithelial migration, we performed bulk RNA-sequencing (RNA-seq) on organoid samples at the start (0 hours), early (36 hours) and active migration (72 hours) stages. Using Gene Set Enrichment Analysis (GSEA), we compared the 36 hour- or 72 hour-sample with the 0 hour-sample and found that gene expression in the EMT pathway ^21^ was not significantly changed in either case (Figure 2A, B). When genes in the EMT pathway were individually examined (Figure 2C), we again found that only *Snail1* and *N-Cad* expression increased, while *Pdgfrb* expression decreased with no significant changes in expression of other EMT markers, which is consistent with the above earlier results.

**Figure 2:**
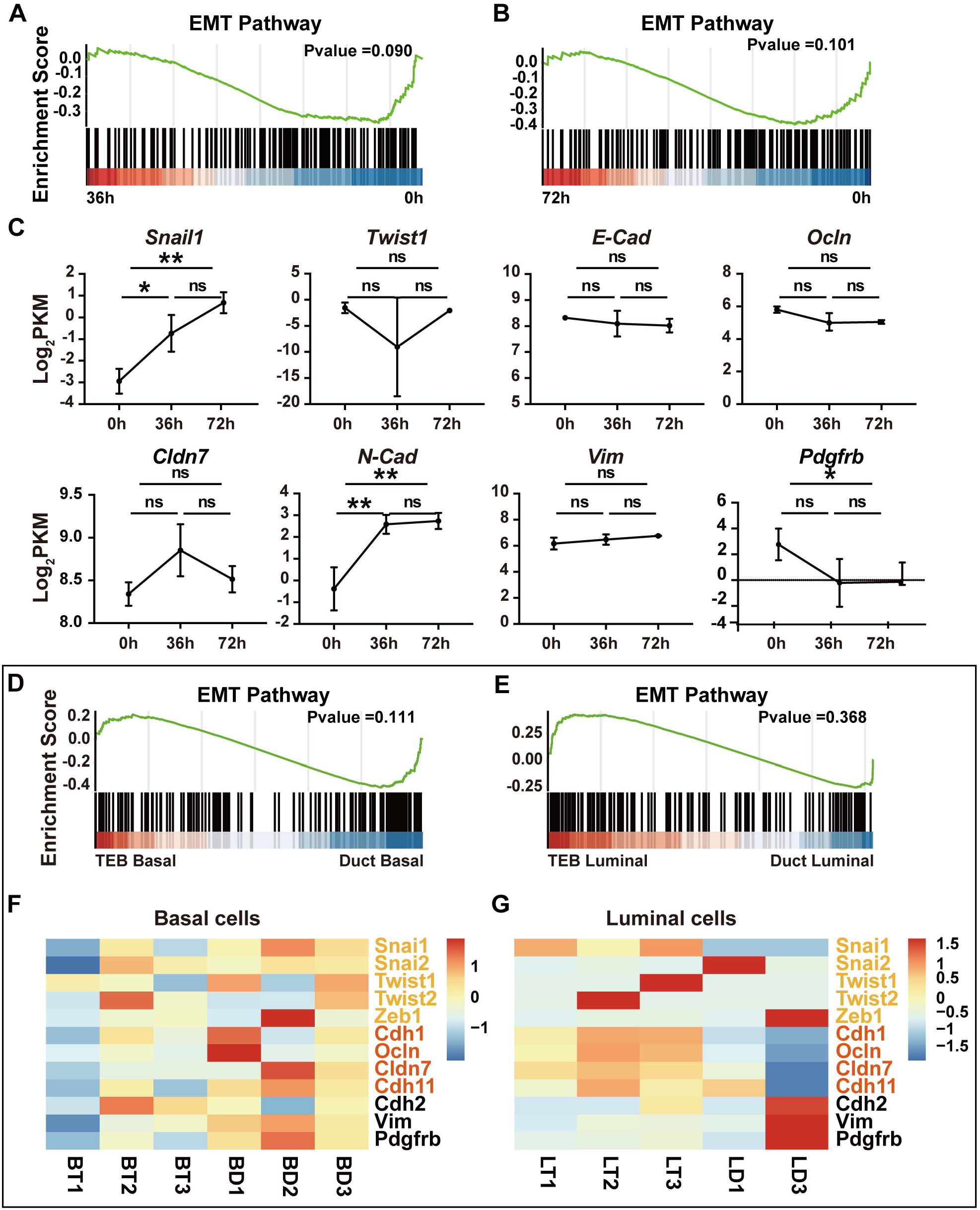
Mammary epithelium does not undergo EMT at the transcriptomic level. (**A, B**) GSEA analyses of EMT pathway between organoids at 36 hours and 0 hours (A) and 72 hours and 0 hours (B) based on bulk RNA-seq of the transcriptomes of organoids at the corresponding stages. Note the P values were not significant. (**C**) Relative mRNA expression of select EMT marker genes at the indicated stages of mammary organoid migration. (**D-G**) GSEA analyses (**D**, **E**) and heatmap analyses of select members (**F**, **G**) of EMT pathway between basal cells (**D, F**) or luminal cells (**E, G**) in the TEBs and the ducts of the branching mammary epithelium at the 5-week stage. RNA-sequencing data were derived from GSE164307^22^. Note the P values were not significant. Data are mean ± SD. Statistical analysis was performed using unpaired Student’s t test. *p < 0.05; **p < 0.01; ****p < 0.0001; n.s., not significant.

Our previous results show that the ducts and the TEBs of the in vivo branching mammary epithelium resemble the in vitro stationary and migrating organoids, respectively ^15^. Using published single-cell RNA-sequencing (scRNA-seq) data ^22^, we examined whether EMT pathway is activated in TEB basal or luminal cells when compared with those in the ductal epithelium. We found that EMT pathway was not significantly changed in TEB basal or luminal cells when compared with those in the duct (Figure 2D, E). Heatmap of a select number of EMT marker genes, including *Snail*, *Twist*, *E-Cad*, etc., in TEB or ductal basal and luminal cells, showed that the expression of these genes was not changed as would have been predicted in the TEBs and ducts (Figure 2F, G).

Taken together, our results show that epithelial cells in neither the in vitro migrating organoids nor in vivo branching TEBs activate the EMT pathway during the collective migration process.

### mRNA Expression of Cell-Cell Adhesion Components Does Not Change as Predicted by Classical EMT Models

A central tenet underlying EMT models is that cell-cell adhesion is reduced during the transition. Therefore, we examined the expression of components of all major cell-cell junctions. We found that expression of most junctional components, including adherens junctions, gap junctions, and hemidesmosomes—the latter being responsible for cell-matrix adhesion— did not significantly change during epithelial migration, as indicated by GSEA analyses (Figure 3A, B, Supplementary Fig. 2A, B, D, E). However, a few components of theses junctions, for example, *Eppk1* of hemidesmosomes, showed decreased mRNA expression during migration, as would be predicted by EMT models (Supplementary Fig. 2F). By contrast, many components of the junctions, including *Ctnna1* (encoding a-catenin) of AJs (Figure 3C), *Gja1*, *Gjb3*, and *Panx1* of gap junctions (Supplementary Fig. 2C), and *Col17a1*, *Lama3*, etc., of hemidesmosomes (Supplementary Fig. 2F) showed increased mRNA expression during epithelial migration, which would be opposite to the predictions from classical EMT models.

**Figure 3:**
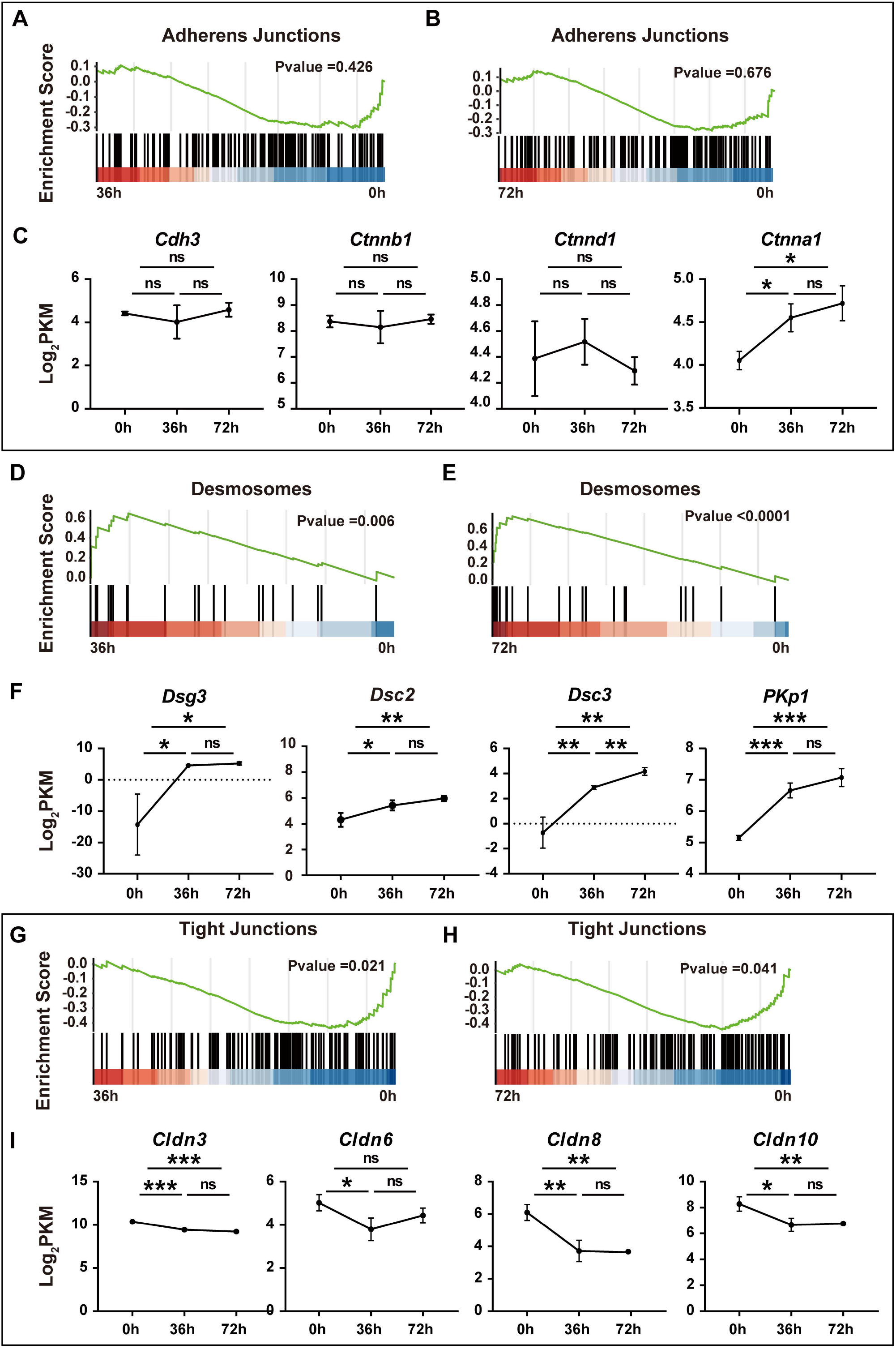
mRNA Expression of Cell-Cell Adhesion Components Does Not Change as Predicted by Classical EMT Models. (**A, B**) GSEA analyses of adherence junction genes between organoids at 36 hours and 0 hours (**A**) and 72 hours and 0 hours (**B**). Note the P values were not significant. (**C**) Relative mRNA expression of select adherence junction genes at the indicated stages of mammary organoid migration. (**D, E**) GSEA analyses of desmosomes genes between organoids at 36 hours and 0 hours (**D**) and 72 hours and 0 hours (**E**). Note the P values were significant. **(F)** Relative mRNA expression of desmosomes genes at the indicated stages of mammary organoid migration. (**G, H**) GSEA analyses of TJ genes between organoids at 36 hours and 0 hours (**G**) and 72 hours and 0 hours (**H**). Note the P values were significant. **(I)** Relative mRNA expression of select TJ genes at the indicated stages of mammary organoid migration. Data are mean ± SD. Statistical analysis was performed using unpaired Student’s t test. *p < 0.05; **p < 0.01; ***p < 0.001; ****p < 0.0001; n.s., not significant.

Interestingly, the desmosome pathway was increased in the migrating epithelium at both the 36 hour-stage and the 72 hour-stage when compared to the stationary epithelium at 0 hours (Figure 3D, E). A closer look at the expression of the individual desmosome components, including *Dsg2*, *Dsg3*, *Dsg4*, etc., show that they had increased mRNA expression over the course of epithelial migration (Figure 3F, Supplementary Fig. 3A).

The only cell-cell junction that was significantly downregulated during epithelial migration according to the prediction based on the classical EMT model was the tight junction (TJ) pathway (Figure 3G, H). An examination of the individual TJ components including *Cldn3*, *Cldn6*, *Cldn8*, etc., further confirmed decreased mRNA expression during epithelial migration (Figure 3I, Supplementary Fig. 4B).

Together, the results show that, except for the TJ pathway, most cell-cell junction pathways are not reduced during epithelial migration as predicted by the pathway analysis.

### Tight Junctions Are a Barrier to, But Cell-Cell Adhesion is Required for, Epithelial Migration

The reduction of TJ component expression suggests that TJs are a barrier to epithelial migration. Therefore, we used both loss and gain of TJ experiments to test this possibility. For the loss of TJ experiment, we aggregated dissociated mammary epithelial cells and subjected the aggregates to the migration assay before TJs could be formed in the Matrigel, as indicated by a lack of band-like staining by ZO1 immunofluorescence (Supplementary Fig. 4A). As we previously documented, mammary organoids showed a ∼30 hour-lag phase before they started to migrate (Figure 4A, C). By contrast, aggregates from dissociated epithelial cells started to migrate much sooner than organoids (Figure 4B, C), suggesting that TJ removal promotes epithelial migration.

**Figure 4:**
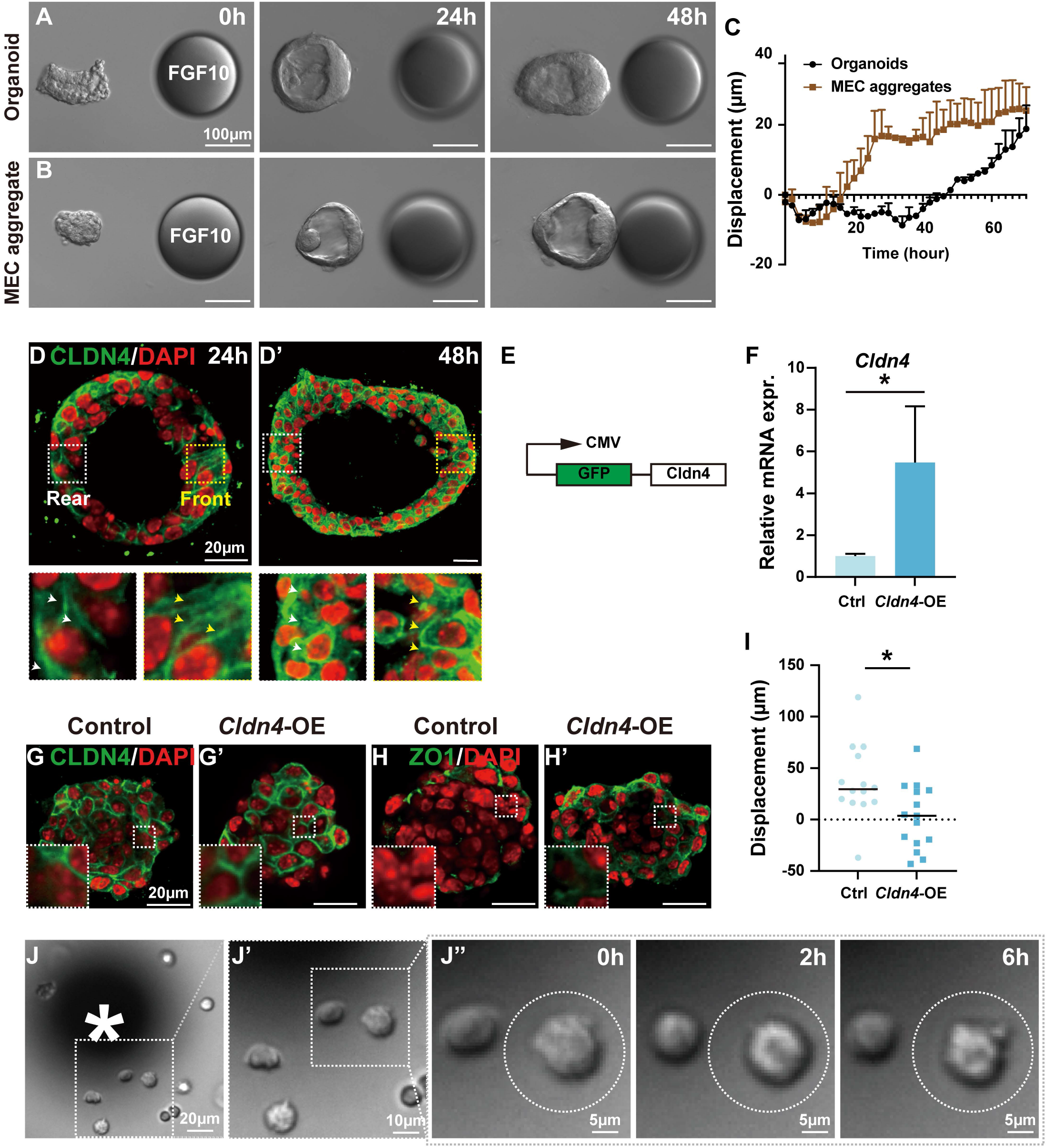
Tight junctions are a barrier, but cell-cell adhesion is required for epithelial migration. (**A-C**) Migration assays of mammary epithelial organoids (**A**) or cell clusters formed after aggregating overnight from mixtures of single basal and luminal cells (**B**), respectively. Quantification of the displacement of organoid and basal-luminal aggregates during the assay duration. X-axis is time (hours) (**C**). N= 2. Scale bars, 100 μm. (**D**) Protein expression and localization of CLDN4 as detected by Immunofluorescent staining at different stages of epithelial migration. Insets with yellow and white dash-lines indicate areas with closed-up views at the front and rear, respectively, of the organoid. White and yellow arrowheads point to the cytoplasmic/membranous localization of the protein assayed. N=3. Scale bars: 20 μm. (**E-H**) Effect of *Cldn4* overexpression on TJs. (**E**) Design of the lentiviral overexpression construct. (**F**) Relative mRNA expression of *Cldn4* using lentivirus carrying either a control construct or an overexpressing vector in mammary epithelial cells. (**G**-**H**) Immunofluorescence of CLDN4 (**G**, **G’**) and ZO1 (**H**, **H’**) on MEC aggregates transfected with either a control or experimental lentivirus expressing GFP alone (**G, H**) or a fusion GFP-CLDN4 protein (**G’, H’**). Note that GFP is membranous in the experimental group due to fusion to CLDN4 (**G’**). Also note that ZO1 expression is increased in *Cldn4*-OE organoids, although the presence of microlumen is not obvious (**H’**). Scale bars, 20 μm. (**I**) Total displacement as calculated after 72 hours using the migration assay. n = 14 for control organoids, and n=15 for *Cldn4*-OE organoids. **(J)** DIC images of the time course of single MECs in adjacent to a FGF10 bead (* asterisk). Note that while the cells were alive, changed shape, and spined, they did not migrate toward the bead (J’-J’’). n=41 cells. Data are mean ± SD. Statistical analysis was performed using unpaired Student’s t test. *p < 0.05. Abbreviations: MEC, mammary epithelial cell.

Previous reports show that forceful expression of claudin causes ectopic TJ formation in fibroblasts ^23^. Therefore, we next examined whether overexpressing *Cldn4* inhibits mammary gland epithelial migration. Using immunofluorescent confocal microscopy, we confirmed that CLDN4 protein localizes to the luminal epithelium as expected (Supplementary Fig. 1D; Figure 4D, D’). To overexpress *Cldn4*, we cloned its coding region, under the control of a strong CMV promoter, into a lentiviral expression vector (Figure 4E). Using qPCR, we detected a five-fold increase of *Cldn4* mRNA expression in MECs transfected with a *Cldn4-*OE construct when compared to MECs transfected with a control vector (Figure 4F).

Using immunofluorescence, we confirmed that CLDN4 is overexpressed in MECs transfected by the *Cldn4-*OE construct, and that overexpressed CLDN4 correctly localized to the plasma membrane (Figure 4G, G’). CLDN4 overexpression caused increased ZO1 expression when compared with control (Figure 4H, H’). However, we did not observe a lumen with a stereotypical belt shaped ZO1 expressionindicative of TJ formation (Figure 4H, H’). The data suggest that, while CLDN4-OE could increase the expression of TJ components, it is insufficient to cause ectopic TJ formation on its own. Interestingly, when subjected to the in vitro migration assay, MECs overexpressing *Cldn4* showed a significant deficit migrating toward the FGF10- bead when compared to control (Figure 4I; Supplementary Fig. 4B, C), thus supporting the prediction that increased TJ expression prevents epithelial migration.

To determine whether cell-cell adhesion overall is a migration barrier, as is currently thought to be the case with EMT, we dissociated mammary organoids and subjected single MECs directly to the migration assay. Interestingly, we found that single MECs, which would typically be defined as individual mesenchymal cells, were unable to migrate toward the FGF10- bead, despite being alive, as indicated by their constant shape-change or spinning movement during the migration assay (Figure 4J-J’’; Movie 1).

Together, the data show that TJs are a barrier to mammary epithelial migration. However, contrary to classical views of EMT, which propose that cell-cell adhesion overall is inhibitory to migration, we show that epithelial cells need to be in contact with each other and exhibit cell adhesion to directionally migrate. These data, and our previously published work ^15^, are thus largely inconsistent with classical EMT models.

### *Snail1* Regulates Moving Speed and Direction of Individual Cells During Epithelial Migration

Based on these findings, we hypothesized that mammary epithelial migration occurs independently of EMT. Consequently, we predicted that this migration would not rely on the classical EMT transcription factor *Snail1*. To test this, we employed a lentivirus carrying shRNA targeting *Snail1* to knock down its expression. Compared to control cells expressing scrambled shRNA, *Snail1* expression was reduced by approximately 70% in the *Snail1*-KD cells (Supplementary Fig. 5A). Contrary to our expectations, we found that *Snail1-*KD aggregates were completely unable to migrate, while control aggregates migrated toward the FGF10-bead over the three-day assay (Figure 5A, B). The inability of *Snail1*-KD cells to undergo epithelial migration suggests that *Snail1* function is essential for the process.

**Figure 5:**
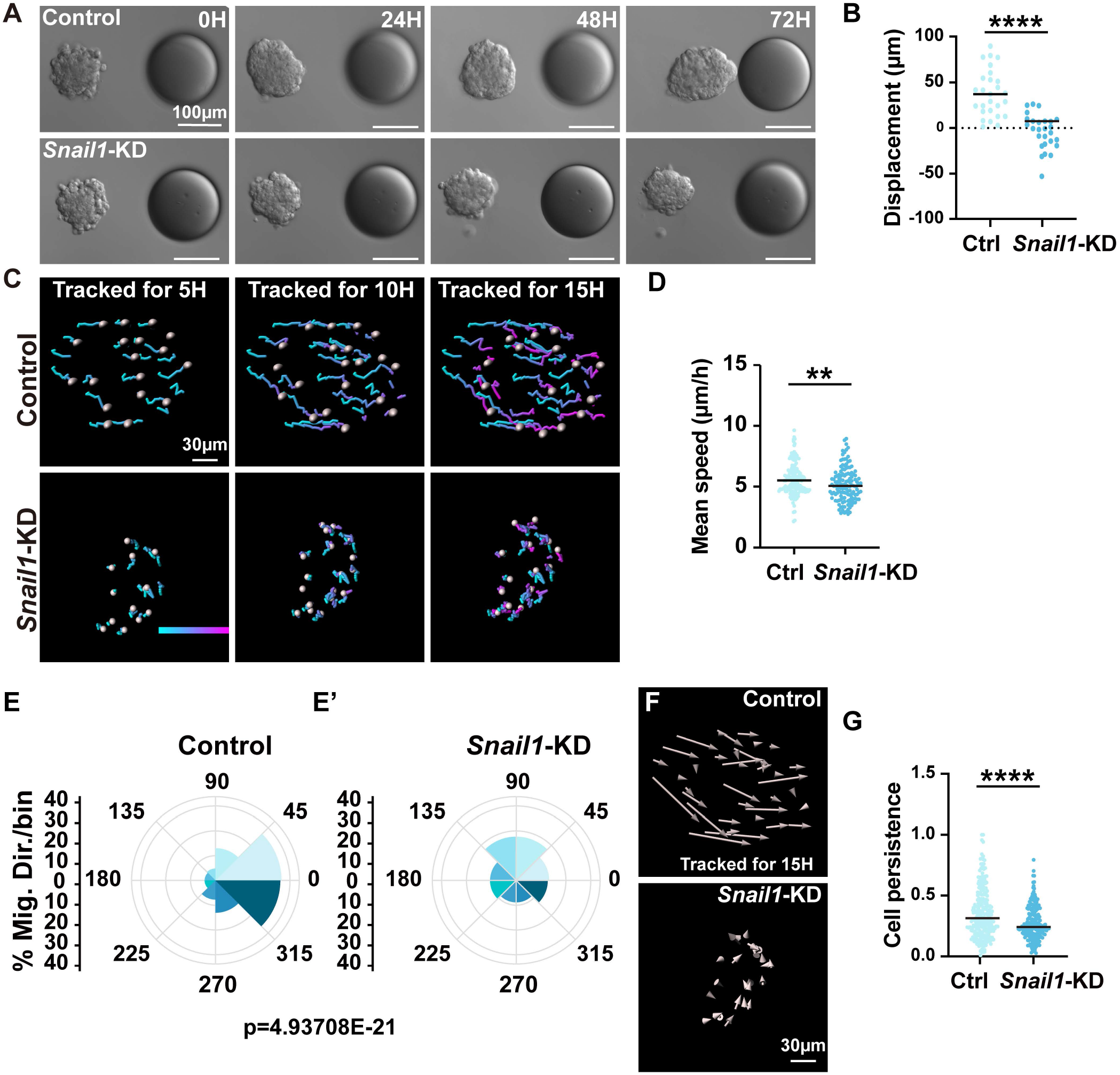
*Snail1* regulates moving speed and direction of individual cells during epithelial migration. (**A**, **B**) Effect of *Snail1* knockdown (KD) using a lentiviral construct expressing either a scrambled RNA or one that targets the gene in the HC11 cell line. Time course (**A**) and (**B**) quantification of displacement of migration of the control (n = 27) or *Snail1-*KD (n = 27) HC11 aggregates toward beads pre-soaked in FGF10. (**C**-**G**) Migration analyses, including cell tracks (**C**), mean speed (**D**), migration direction (**E, E’**), and persistence (**F**, **G**) of individual cells from the control (n of aggregates = 7, n of tracks =259) and *Snail1*-KD (n of aggregates = 7, n of tracks =200) aggregates. Note that movement angle is defined as the angle of movement direction and that toward the FGF10 bead. For example, the angle equals to zero when a cell moves directly toward the FGF10 bead (**E**, **E’**). Also note that cell persistence is defined as a cell’s tendency to maintain its direction of movement over time. Data are mean ± SD. Statistical analysis was performed using unpaired Student’s t test. **p < 0.01; ****p < 0.0001; N.S., not significant. Abbreviations: Ctrl, control; Mig, migration; Dir, direction.

To analyze the migratory defects in *Snail1*-KD cells, we used a mosaic analysis to track individual cells during the migration process. Specifically, we mixed control or *Snail1*-KD mCherry-expressing cells with GFP-expressing control or *Snail1*-KD cells, respectively, at a 1:9 ratio, and then tracked the movement of red cells during migration (Figure 5C; Movies2, 3). The mean speed of *Snail1*-KD cells dropped ∼10% when compared to control cells (Figure 5D), and in contrast to control cells, which all migrated toward the FGF10 signal, the migration direction of *Snail1*-KD cells was randomized (Figure 5E, E’). Likewise, cell persistence, which measures the tendency of a cell to continue migrating in the same direction calculated as track displacement divided by total track length, was significantly reduced in *Snail1*-KD cells when compared to control cells (Figure 5F, G).

Together, these results show that *Snail1* plays an essential role in epithelial migration by regulating movement speed, and more importantly direction, of individual cells during collective migration. In the absence of *Snail1*, movement direction of individual mutant cells was randomized, and, as a result, mutant cell clusters were unable to undergo directional migration.

### *Snail1* Promotes Cell-Cell Adhesion and Multiple Aspects of Directional Migration

To determine the transcriptional role of *Snail1*, we used RNA-sequencing and compared the transcriptomes of *Snail1*-KD cells and controls during epithelial migration. Specifically, we subjected *Snail1*-KD cells and control cells to the migration assay and prepared samples for bulk RNA-seq at the 36-hour stage of migration. Surprisingly, GSEAs showed that both the tight and adherens junctions pathways were significantly reduced in *Snail1*-KD cells when compared to control cells (Figure 6A, B). This is contrary to the generally held notion that *Snail1* negatively regulates adherence and TJs components at the transcriptional level ^24,25^, which would have predicted an expression increase of these pathways. When examined more closely, however, most of the individual genes of the main TJ components were not significantly changed the transcriptional level (Supplementary Fig. 5B). This was confirmed by our qPCR results of the same TJ component genes (Supplementary Fig. 5C). Likewise, we found that the individual genes of the main adherens junctions components were not significantly changed at the transcriptional level (Supplementary Fig. 5D). Although an increased level of mRNA expression was not observed, the results were nonetheless surprising because they confirmed that, contrary to the common belief, *Snail1* inhibits the transcription of neither the tight nor adherens junctions pathway.

**Figure 6:**
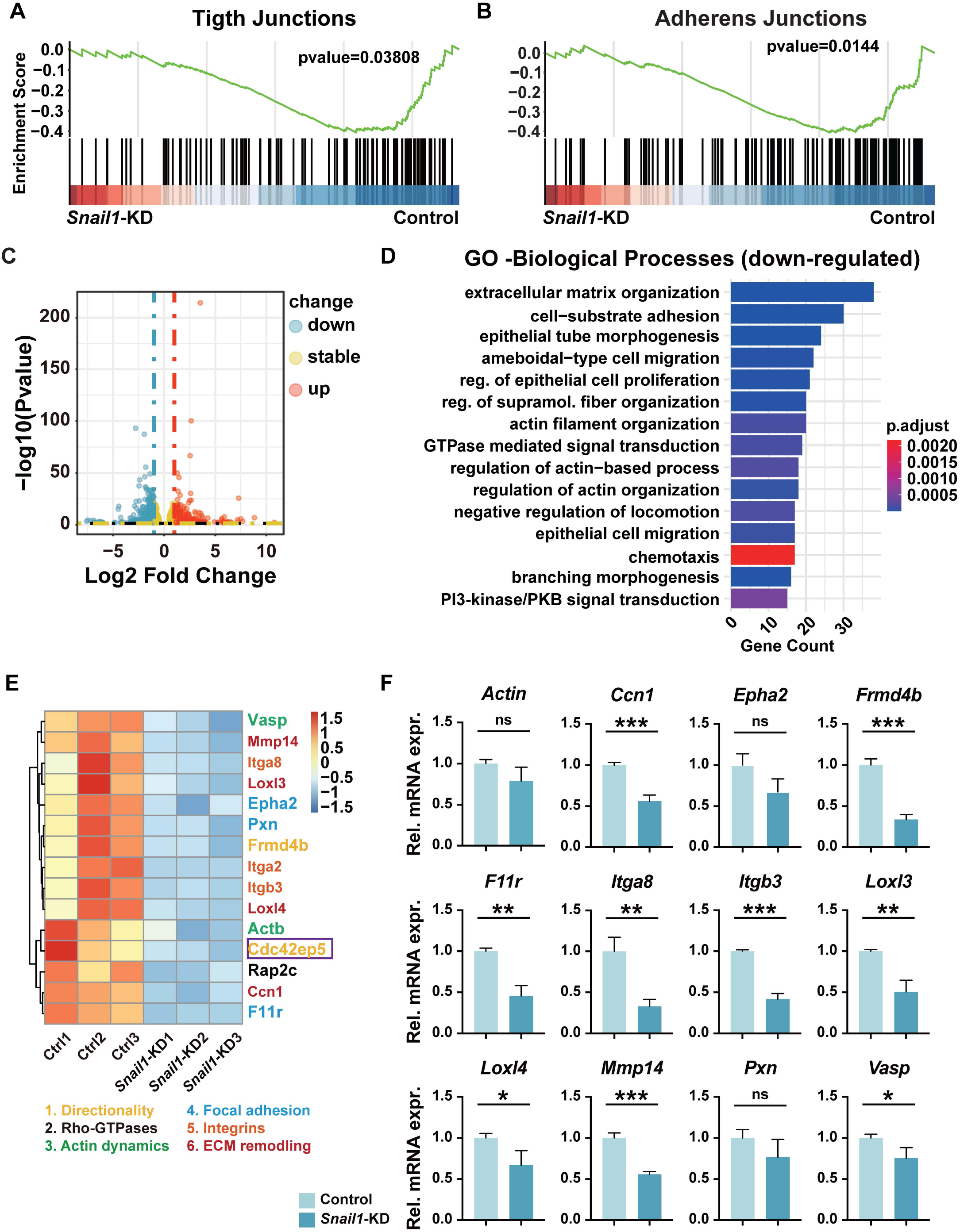
*Snail1* promotes cell-cell adhesion and multiple aspects of directional migration. (**A**, **B**) GSEA analyses of the TJs pathway (**A**) and the adherence junctions pathway (**B**) between *Snail1-*KD and control aggregates. 3D migration assay using HC11 cells infected with either control or *Snail1*-KD virus. Samples were collected at 36 hours post-experiment for sequencing, followed by GSEA or expression analysis. Note the P values were significant. **(C)** Volcano plot based on the top 100 most highly down-and up-regulated genes in *Snail1*-KD HC11 aggregates when compared with control aggregates. **(D)** Gene ontology analysis of the differentially expressed genes (p < 0.05) between *Snail1*-KD HC11 aggregates and control aggregates to determine the biological processes with which these genes might be involved. (**E**, **F**) Heatmap (**E**) and qPCR validation (**F**) of select candidate regulators of cell migration from the top 100 most highly down-regulated genes. Note that candidate genes are categorized into groups with presumed functions in the sequential steps of directional cell migration ^18^. Data are mean ± SD. Statistical analysis was performed using unpaired Student’s t test. *p < 0.05; **p < 0.01; ***p < 0.001; ****p < 0.0001; N.S., not significant. Abbreviations: Ad, adenovirus; Ba, basal; Lu, luminal; TEB, terminal end buds.

As shown by Volcano plot, expression of many essential genes was impacted in *Snail1*-KD cells (Figure 6C). GO analysis of the down-regulated genes further showed that the main biological processes affected include ECM organization, cell substrate adhesion, actin filament organization, chemotaxis, GTPase-mediated signal transduction, PI3 Kinase signaling, etc. (Figure 6D), almost all of which are known to be essential for directional migration. Additionally, many genes were found to be upregulated in *Snail1*-KD cells. GO analysis indicated that these genes are involved in the regulation of viral processes, viral defense response, and antiviral innate immune responses (Supplementary Fig. 6E). The implications of these findings— particularly the unexpected role of *Snail1* in repressing immune functions—remain unclear in the context of its role in migration.

These data thus suggest that *Snail1* regulates multiple aspects of epithelial migration. To validate this possibility, we narrowed the list of the potential *Snail1* targets down to 15 genes encompassing the main sequential steps of directional cell migration, including directionality setup, Rho-GTPase activation, actin dynamic regulation, etc. (Figure 6E). Using qPCR reactions, we confirmed that most of these candidate genes had significantly reduced mRNA expression in *Snail1*-KD cells compared to control cells, except for *Pxn*, *Actin*, and *Vasp*, which showed a trend toward reduced expression, but this was not significant (Figure 6F).

Together, these results support the notion that *Snail1* regulates multiple aspects of directional migration. However, it does not repress adherens or TJs at the transcriptional level as commonly believed.

### *Cdc42ep5* Regulates Moving Direction During Epithelial Migration

Of the potential *Snail1* target candidates that regulate directionality, *Cdc42ep5* (*Ep5*) was chosen for further analysis because it encodes an effector of *Cdc42*, a known regulator of front-rear polarity and directional migration ^26^. We found that *Ep5* was significantly downregulated in *Snail1-*KD cells based on the bulk RNA-seq data (Figure 7A), suggesting that *Ep5* is a *Snail1* target gene. Based on published scRNA-seq datasets, we found that *Ep5* is expressed in both basal and luminal cells, though more highly in the latter throughout development, including the 5-week, 10-week, and pregnancy day 12 (P12; Figure 7B). To assess the impact of Ep5 knockdown on the directional migration of HC11 cells, we generated a lentiviral construct carrying an shRNA sequence targeting Ep5. This reduced Ep5 mRNA expression by 70% in HC11 cells compared to a control lentivirus expressing a scrambled shRNA sequence (Figure 7C). When subjected to the migration assay, *Ep5* aggregates showed a significantly reduced ability to migrate compared to the control aggregates (Figure 7D, E; Supplementary Fig. 6A).

**Figure 7:**
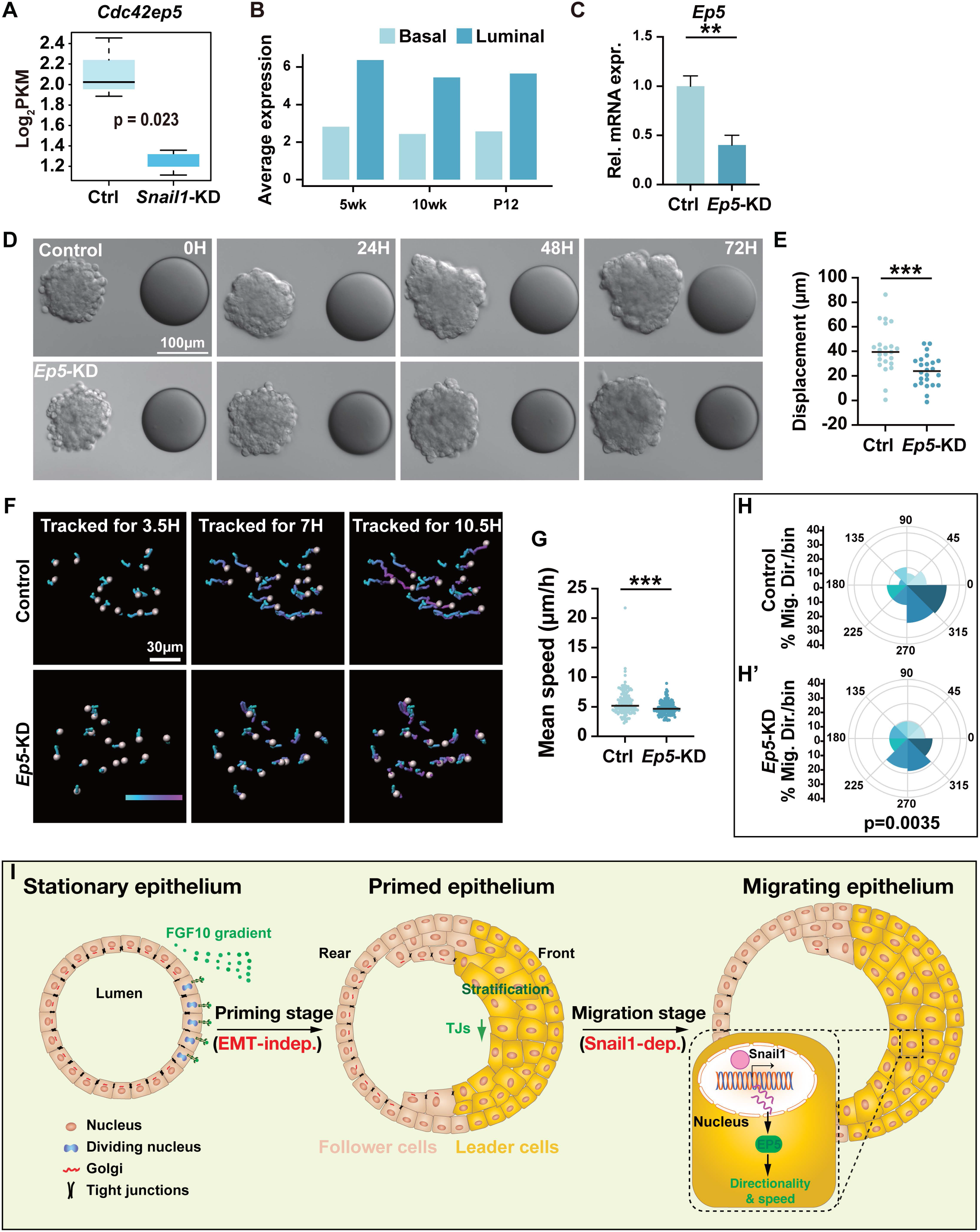
*Cdc42ep5* regulates moving direction during epithelial migration. **(A)** Comparison of expression levels between the control and *Snail1*-KD migration samples at 36 hours in *Ep5*. **(B)** Expression levels of *Ep5* in basal and luminal cells at different developmental stages, based on single-cell data analysis (access # GSE164017). (**C**-**E**) Effect of *Cdc42ep5* (*Ep5*) knockdown (KD) using a lentiviral construct expressing either a scrambled RNA or one that targets the gene in the HC11 cell line. (**C**) knockdown efficiency as measured by qPCR. Time course (**D**)and quantification of displacement (**E**) of epithelial migration of the control (n = 23) or *Ep5*-KD (n = 24) HC11 aggregates toward beads pre-soaked in FGF10. (**F**-**H’**) Migration analyses, including cell tracks (**F**), mean speed (**G**), migration direction (**H, H’**), of individual cells from the control (n of aggregates = 6, n of tracks =139) and *Ep5*-KD (n of aggregates = 5, n of tracks =139) aggregates. Movement angle is defined as the angle of movement direction and that toward the FGF10 bead. Data are mean ± SD. Statistical analysis was performed using unpaired Student’s t test. *p < 0.05; **p < 0.01; ***p < 0.001; ****p < 0.0001; N.S., not significant. See also. (**I**) Model diagram of EMT-independent mammary epithelial migration. Based on our data reported here and previously, we propose that mammary epithelium can change from a stationary to a migratory state via a two-step processes. In the first step, the chemotactic signal FGF10 promotes cell proliferation and stratification in the organoid front. This leads to a reduction of TJs and conversion of apicobasal polarity to front-rear polarity. As a result, leader cells are generated, and epithelium becomes “primed,” ready to enter the next phase. This step is independent of EMT. In the second step, leader cells, which are a dynamic cell population that forms intraepithelial protrusions, start to migrate directionally toward FGF10. We show that *Snail1* is essential for this migration process by transcriptionally regulate *Ep5*, a cofactor of the front-rear polarity regulator *Cdc42*. In addition to directionality, *Snail1/Ep5* also regulates other aspects, for example, speed of individual cells during collective migration. Mammary epithelium maintains epithelial features throughout the course of migration. Abbreviations: indep, independent; dep, dependent; TJs, tight junctions.

Using a mosaic analysis similar to the one mentioned above, we tracked individual cell movement during directional migration of *Ep5* and control aggregates. The mean speed of *Ep5*- KD cells dropped ∼10% when compared to control cells (Figure 7F, G; Movies 4, 5). Importantly, when compared to control cells, which migrated toward the FGF10 signal, *Ep5*-KD cells showed a random directional preference (Figure 7H, H’). Likewise, cell persistence was significantly reduced in *Ep5*-KD cells when compared to control cells (Supplementary Fig. 6B- C).

These results show that *Snail1* regulates directional migration of mammary epithelium by targeting *Ep5* transcription.

## DISCUSSION

Traditionally, epithelium has been considered immobile, requiring EMT for migration, either individually or collectively as mesenchymal cells. Building on our previous studies demonstrating unique mammary epithelial migration from mesenchymal collectives, we now conclusively show that EMT is not required for epithelial migration in this tissue. This conclusion is supported by cellular and molecular evidence, including EMT marker expression at the mRNA, protein, localization, and transcriptomic levels. Although TJs serve as barriers to, but cell-cell adhesion is essential for, epithelial migration. Moreover, contrary to long-held assumptions, *Snail1* does not repress cell-cell adhesion. Instead, it plays a crucial role in epithelial migration by regulating the directionality of individual cells via *Cdc42ep5* (Figure 7I).

### Mammary Epithelium Does Not Transition to A Mesenchymal State for Migration

Our investigation, which examined both commonly used EMT markers and associated molecular events, definitively demonstrates that mammary epithelium does not undergo EMT before migration. Specifically, most EMT markers do not show the expected changes in mRNA or protein expression, or localization, during migration. Additionally, despite previous suggestions^27^, the EMT pathway is not upregulated during in vitro migration or in the TEBs.

At the molecular level, we found that TGF-β, a well-known EMT promoter, not only fails to induce mammary epithelial migration but also strongly inhibits it. Similarly, *Snail1*, one of the most extensively studied EMT transcription factors, does not repress cell-cell adhesion, as is commonly assumed. Interestingly, our data suggest that *Snail1* is required for directing individual cell movements during active epithelial migration.

With respect to cell-cell adhesion, which traditionally distinguishes epithelium from mesenchyme, we observed that most adhesion types, particularly adherens junctions, are not downregulated during migration. In fact, desmosomes are upregulated. The only cell-cell adhesion structure that decreases, consistent with EMT predictions, is the TJs. Our gain- and loss-of-function studies demonstrate that TJs serve as barriers to epithelial migration.

The requirement for TJ downregulation aligns with our previous findings that, during the stationary phase of mammary epithelium, the leading front of epithelial cells undergo active proliferation, stratification, and conversion from apicobasal polarity to front-rear polarity ^15^. These sequential events generate a dynamic population of leader cells necessary for the directional migration of the mammary epithelium ^18^. Together, these findings challenge long-standing beliefs and suggest that mammary epithelium can migrate without undergoing EMT (Figure 7I).

### Does Mammary Epithelium Undergo a “Partial” EMT?

Several studies have reported that, in various developmental or cancer contexts, only a subset of EMT markers show the expected changes ^9–11^. This has led to the concept of “partial” EMT, a hybrid state between epithelial and mesenchymal phases ^5,12^. Given that TJs are reduced as expected, and some EMT markers change in line with EMT models, could mammary epithelium undergo a partial EMT and enter a hybrid epithelial-mesenchymal state before migration?

The answer depends on how we define epithelium and mesenchyme, which is central to determining what constitutes a "full" or "partial" EMT. The definitions of both epithelium and mesenchyme have evolved over time, reflected in the growing number of behavioral, structural, and molecular markers used to describe these tissues. Therefore, it is essential to evaluate the accuracy—or lack thereof—of these markers in the context of their original definitions: epithelium as cell sheets lining body surfaces and cavities, and mesenchyme as epithelial-derived loose cells embedded in the extracellular matrix^28,29^. These definitions influence our interpretation of whether epithelial tissue transitions into a mesenchymal state.

Due to the dominance of classical views of EMT, epithelium and mesenchyme are often equated with stationary and migratory states, respectively, and loss of apicobasal polarity is commonly considered a hallmark of EMT^4,5^. However, our data, both here and previously ^15,18,19^, suggest that mammary epithelium, like mesenchyme, can exist in either a stationary or migratory state, transitioning between these phases depending on the context. A loss of apicobasal polarity to gain front-rear polarity, or directionality, is part of this epithelial transition from a stationary to migratory state (Figure 7I).

Our data also reveal that EMT markers are differentially expressed in subtypes of stationary mammary epithelium. For example, "epithelial" markers such as *E-Cad*, *Ocln*, and *Cldn4* are restricted to the luminal epithelium, while "mesenchymal" markers like *Twist1*, *N-Cad*, *Vim*, and *Pdgfrb* are preferentially expressed in the basal epithelium, which is still classified as epithelial based on its anatomical and morphological characteristics. Importantly, this differential expression pattern of EMT markers persists throughout various stages of mammary epithelial migration, arguing against their use as definitive markers of EMT.

Thus, we conclude that current EMT markers are too subjective to accurately reflect the mechanisms underlying epithelial transitions from stationary to migratory states. Until more precise markers are established, it is premature to describe these processes as "partial" or "full" EMT events. Instead, we propose that mammary epithelium retains epithelial characteristics throughout the stages of directional migration, without transitioning to a mesenchymal state during priming or migration (Figure 7I).

### Reexamining EMT processes

In addition to the issues raised concerning EMT markers, the current understanding of core EMT events appears problematic. For instance, we show that TGF-β inhibits mammary epithelial migration and that *Snail1* does not repress cell-cell adhesion. While these findings may be explained by differences in developmental or pathological contexts, other observations, such as the persistence of most forms of cell-cell adhesion—except for TJs—are harder to reconcile with EMT models. Single epithelial cells, which lack cell-cell adhesion and would traditionally be considered mesenchymal, are unable to migrate. This is consistent with reports that E-Cad is essential for cancer cell metastasis and mesenchymal cell migration ^7,8^.

Despite early studies showing *Snail* function is essential for EMT, for example by repressing cell-cell adhesion including E-Cad expression ^24,25^, neural crest formation is normal in mice lacking both *Snail1* and *Snail2* ^30^. Our data are thus consistent with these in vivo studies suggesting that *Snail1* is not required for EMT. Although *Snail1* does not repress cell adhesion, we show that it is essential for directing mammary epithelial migration by controlling individual cell movements. This suggests that currently presumed EMT regulators may instead function by directing epithelial migration, rather than mediating an EMT transition, as commonly thought.

In summary, we call for a reevaluation of classical EMT models in both developmental and cancer biology. Recognizing that epithelium can migrate without transitioning into a mesenchymal state may help refocus attention on contexts where EMT truly occurs, such as when epithelial cells disseminate from their primary site during processes like formation of mesoderm and neural crest, or cancer metastasis. This reevaluation may lead to a more accurate understanding of developmental processes and cancer progression.

## MATERIALS AND METHODS

### Mouse Strains

Wild-type C57BL/6J mice were obtained from GemPharmatech Co., Ltd. Mice were housed and maintained according to the guidelines of the University of South China’s Institutional Animal Care and Use Committee.

### Preparation of Mammary Organoids and Primary Epithelial Cells

Mammary organoids and primary epithelial cells were extracted using previously described methods^31,32^. Briefly, mammary glands No. 2, No. 3, and No. 4 from mice were collected, minced, and digested in 10 ml DMEM/F12 containing 2 mg/ml collagenase (Sigma, C5138), 2 mg/ml trypsin (Gibco, 27250018), 5 µg/ml insulin (Yeasen, 40107ES25), 5% fetal bovine serum (FBS) (Gibco, 10099141), and 50 µg/ml gentamicin. The mixture was shaken for 25 minutes, centrifuged at 560g for 10 minutes, and the precipitate was collected. Mammary organoids were purified by differential centrifugation at 450g and then digested in TE for approximately 12 minutes to obtain single cells.

### In Vitro 3D Epithelial Migration Assay

Primary mammary organoids or MECs, as well as HC11 aggregates, were used for migration assays. For obtaining MECs aggregates, mammary epithelial cells were digested into single cells, and 2000 cells were sorted via flow cytometry and added to ultra-low attachment 96-well plates (Qingdao Ama Co., Ltd, 16196-6SULH) containing 100 µl of MaSC medium (DMEM/F12 containing 5% FBS, 10 ng/ml EGF [Novoprotein, C029], 20 ng/ml FGF2 [GenScript, Z03116], 4 µg/ml Heparin, 5 µM Y-27632, 1% L-Alu-Gln [Beyotime, C0211], and 1% P/S). The plates were incubated overnight at 37°C to form aggregates. For HC11 cells, 250 cells were added to ultra-low attachment 96-well plates containing 100 µl of assay medium (RPMI 1640 containing 2% FBS, 1% P/S, 1% L-Alu-Gln, 10 ng/ml EGF, 5 µg/ml insulin) and incubated overnight at 37°C for culturing.

For the migration assay, as described before, Heparin sulfate beads were soaked in 10 µl of 100 µg/ml FGF10 (GenScript, Z03155). Then, 80% Matrigel (Corning, 354230) was added to a glass-bottom 24-well plate (Cellvis, P24-1.5H-N) or 8-well chamber (Cellvis, C8-1.5H-N), and beads and organoids were placed approximately 100 µm apart and covered with basic medium (DMEM/F12 or RPMI 1640, 1% P/S, 1% L-Ala-Gln, 1% ITS). Live cell imaging was conducted.

For the experiment of EMT promoter TGF-β protein (Abclonal, RP00671) on FGF10-induced epithelial migration. TGF-β was added to the basic medium at 5ng/ml.

### Cell Culture

HEK293T cells were cultured in DMEM (Gibco, # C12430500BT) supplemented with 10% FBS (Lonsera, S711-001S), 1% sodium pyruvate, 1% non-essential amino acids, 1% L-Ala-Gln, and 1% P/S. HC11 cells were cultured in RPMI 1640 with 10% FBS, 10 ng/ml EGF, 5 µg/ml insulin, 1% L-Ala-Gln, and 1% P/S. Basal and luminal cells were cultured in Advanced DMEM/F12 with 5% FBS, 10 ng/ml EGF, 20 ng/ml FGF2, 4 µg/ml heparin, 5 µM Y-27632, 1% L-Ala-Gln, and 1% P/S.

### Lentiviral Construction, Packaging and Infection

shRNA sequences were designed using BLOCK-iT™ RNAi Designer and cloned into the pLKO.1 vector (Addgene #8453). The vector was digested with AgeI (Abclonal, RK21125) and EcoRI (Abclonal, RK21102), ligated with annealed shRNA fragments using T4 DNA ligase (Abclonal, RK21501), and verified by Sanger sequencing. For overexpression, Claudin4 CDS was cloned into pLeGO-SFFV-P2A-mCherry using the ClonExpress Ultra One-Step Cloning Kit (Vazyme, C115). Plasmids were prepared using an endotoxin-free plasmid extraction kit (Magen, P1112). Knockdown or overexpression efficiency was assessed in HC11 or luminal cells.

HEK293T cells were co-transfected with pMD.2, pspax2, and transfer plasmids using Eztrans (Shanghai Life-iLab Biotech Co., Ltd, AC04L099). Virus-containing medium was collected 60- and 88-hours post-transfection, concentrated by centrifugation at 27,000 rpm for 2 hours, and resuspended in DMEM/F12 with 10% FBS. Mammary organoids or cells were digested into single cells for infection. Lentivirus infection lasted 2 hours in ultra-low attachment culture dishes, and positive cells were sorted by flow cytometry. The shRNA sequences were as follows: Scramble shRNA: 5’-CCTAAGGTTAAGTCGCCCTCG-3’. Snail1-shRNA: 5’-GCATGTCCTTGCTCCACAAGC-3’. Cdc42ep5-shRNA: 5’-GTCCCTACAATCCCTTCAAAG-3’.

### Western Blotting

Primary cells or organoids were lysed in RIPA buffer (Beyotime, P0013B) with 1 mM phenylmethylsulfonyl fluoride, 2 µg/mL leupeptin (Yeasen, 20112ES50), and 10 µg/mL aprotinin (Beyotime, SG2004). After centrifugation at 13,500g for at 4 °C 10 minutes, protein concentrations were determined using a BCA kit (EpiZyme, ZJ102). 5-10 µg of protein was separated via Tris-glycine electrophoresis and transferred to PVDF membranes. After blocking with 5% BSA (Sigma, #WXBC3116V) or skim milk, target proteins were visualized using anti-E-cadherin (Abclonal, A20798), anti-CLDN7 (Invitrogen, 34-9100), anti-OCLN (Invitrogen, 40- 6100), anti-Vimentin (HUABIO, ET1610-39), anti-Actin (Abclonal, AC006). After the primary antibody was incubated and extensively washed, the membrane was reacted with the HRP-conjugated secondary antibody at room temperature for 2h. The reactive bands were developed by ECL kit (EpiZyme, SQ202) and tested with Amersham Imager 680. For the quantification of protein expression, band density was measured using Image J software.

### Immunofluorescence Staining

Organoids or sections fixed with 4% PFA (Beyotime, P0099) for 30min or 15min were permeabilized in 0.5% Triton X-100 for 2 hours at RT, blocked overnight at 4°C in PBS containing 10% goat serum (Meilunbio, MA0233) and 0.2% Tween 20 and then were incubated with a primary antibody at 4 °C overnight. Subsequently, a secondary antibody was applied at RT for 2 hours, and DAPI (Beyotime, C1005) was used for nuclear staining. Confocal imaging was performed using the *Nikon CSU W1*-01 *SoRa*+NIR spinning disk microscope. Primary antibodies used in this study were E-cadherin antibody (Abclonal, A20798), Claudin7 antibody (Invitrogen, 34-9100), Vimentin antibody (HUABIO, ET1610-39), N-cadherin antibody (Abclonal, A19083), Claudin4 antibody (Invitrogen, 36-4800), ZO-1 antibody (Proteintech, 21773-1-AP). See Supplementary Table 1 for antibody information.

### Quantitative Real-Time PCR

RNA was extracted using an RNA extraction kit (Abclonal, RK30120) or Micro-scale RNA extraction kit (Magen, R4125) and reverse transcribed into cDNA using ABScript III RT Master Mix for qPCR (Abclonal, RK20429). qPCR was performed with 2X Universal SYBR Green Fast qPCR Mix (Abclonal, RK21203) on a BIO-RAD CFX Connect Real-Time System. *Actb* was used as the internal reference. Primer sequences are listed in Supplementary Table 2.

### Image Acquisition and Time-Lapse Imaging

Confocal images were acquired using the Nikon CSU W1-01 SoRa+NIR Spinning Disk Microscope with CFI Plan Apochromat VC 20X or CFI Apo LWD 40XWI λS objectives. Time-lapse imaging was performed at 37°C with 5% CO, capturing images every 15 or 30 minutes. DIC (differential interference contrast) images were acquired using the Zeiss observer z1 microscpoe with LD Plan-Neofluar 20x/0.4 Korr M27 objective. Time course imaging was performed at 24h,48h and 72h respectively. Images were processed with ImageJ software.

### Tracking of Organoids and Cells

ImageJ was used to measure organoid size, distance traveled, and speed, based on the center of gravity of the organoids. Red fluorescent-labeled cells were tracked automatically using the Spots function of Imaris software (Bitplane). The center of each cell was marked with a spot in each frame, and spots were connected over time. Mean cell speed was calculated by dividing the total track length by the tracking time, while displacement length represented the total cell displacement. Cell persistence is defined as the ratio of track displacement to track length, calculated as: cell persistence = track displacement/track length. This formula reflects the directionality of cell movement, with higher values indicating more persistent, linear migration. An unpaired t-test was used to assess the statistical significance of cell speed and displacement lengths between the front and rear cells of the organoids.

### Bulk RNA-Sequencing and Bioinformatics Analysis

For sequencing, HC11 cells infected with scramble or snail1 KD lentivirus were aggregated and subjected to a migration assay. After 36 hours of migration, cells were harvested using Recovery Solution (Corning, #354253) for subsequent RNA extraction. For EMT-related sequencing, conduct in vitro migration assays on the organoids. Collect the migrated samples at 0 hours, 36 hours, and 72 hours for subsequent RNA extraction.

Total RNA was extracted using either the Micro-scale RNA extraction kit (Magen, R4125) or the RNA extraction kit (Abclonal, RK30120) according to the manufacturers’ instructions. RNA quality was assessed using a 5300 Bioanalyzer (Agilent) and quantified with the ND-2000 spectrophotometer (NanoDrop Technologies). Only high-quality RNA samples (OD260/280 = 1.8–2.2, OD260/230 ≥ 2.0, RQN ≥ 6.5, 28S:18S ≥ 1.0, >10ng) were used for sequencing library construction.

RNA purification, reverse transcription, library construction, and sequencing were performed at Shanghai Majorbio Bio-pharm Biotechnology Co., Ltd. (Shanghai, China) according to the manufacturer’s protocols. RNA-seq transcriptome libraries were prepared using the Illumina® Stranded mRNA Prep, Ligation kit (San Diego, CA) with 30 ng of total RNA. Messenger RNA was isolated via poly-A selection using oligo(dT) beads and fragmented using a fragmentation buffer. Double-stranded cDNA was synthesized using the SuperScript double-stranded cDNA synthesis kit (Invitrogen, CA) with random hexamer primers. The cDNA was subjected to end-repair, phosphorylation, and adapter ligation according to library preparation protocols. Libraries were size selected for cDNA fragments of 300 bp using 2% Low Range Ultra Agarose, followed by PCR amplification with Phusion DNA polymerase (NEB) for 15 cycles. Libraries were quantified using a Qubit 4.0 fluorometer, and sequencing was performed on the NovaSeq 6000 platform with the NovaSeq Reagent Kit.

Raw paired-end reads were trimmed and quality-controlled using fastp with default parameters.

Clean reads were aligned to the reference genome using HISAT2 software. Mapped reads were assembled for each sample using StringTie in a reference-based approach.

Raw reads in the fastq format were aligned to the mouse genome (mm10) using the subjunc function of the subread package (Version 2.0.3). Gene counts for each sample were generated from GTF annotation files (gencode.vM30.annotation.gtf) using the FeatureCounts function of the subread package. Differential gene expression was analyzed using the DESeq2 R package, and gene set enrichment analysis (GSEA) was conducted with the clusterProfiler R package. Ligand-receptor interaction data were obtained from (github.com/arc85/celltalker) using the celltalker R package.

### Bioinformatics Analysis of Genes in Different Stages of Mammary Gland Development

scRNA-seq data covering different stages of mammary gland development were obtained from GSE164017. Data input, quality control, integration, and cell clustering were performed using the Seurat R package. Basal cells were marked by Krt14 and Krt5 expression, while luminal cells were identified by Krt18 and Krt8 expression. Gene expression levels were averaged across different developmental stages in both basal and luminal cells.

scRNA-seq and RNAseq data of EMT-related gene expression of basal and luminal cells in TEB, duct of mammary gland come from GSE164017 and GSE164307. Differential gene analysis was performed using DESeq2 and GO enrichment analysis and GSEA were conducted with the clusterProfiler R package.

## Quantification and Statistical Analysis

Sample sizes are indicated in figure legends. Statistical significance was determined using two-tailed Student’s t-tests or one-way ANOVA, or Chi-square test. Error bars represent SD, with significance denoted as ∗p < 0.05, ∗∗p < 0.01, ∗∗∗p < 0.001, and ∗∗∗∗p < 0.0001.

## Data availability

All data supporting the conclusions of the paper are available in the article and corresponding figures. scRNA-seq data were downloaded from public databases as indicated in the methods section.

## ACKNOWLEDGEMENTS

We thank members of the Lu lab for discussions and Drs. Chenleng Cai, and Yikang Rong for thoughtful comments and suggestions. We thank technical support from the Molecular Imaging Core Facility, and Molecular and Cell Biology Core Facility in Hengyang Medical School at South China University. This work was supported by grants from the National Science Foundation of China (32470886 to P.L., 82330035 to K.X. and 32270882 to Y.L.).

## AUTHOR CONTRIBUTIONS

J.C. Data acquisition, analysis, and curation

R.M. Data quality control, analysis, diagrams, and review.

Z.D. Data analysis, bioinformatics analysis.

Y.L., Cldn4 overexpression and its effect on migration. J.C., O.D.K., and K.X., intellectual contributions, review.

P.L., Conceptualization, funding acquisition, data curation, and writing.

