## Supplemental Figure for "Mammary Epithelial Migration is EMT-Independent"

**Sup. Fig1. Mammary epithelium does not show typical EMT signatures during collective migration.**

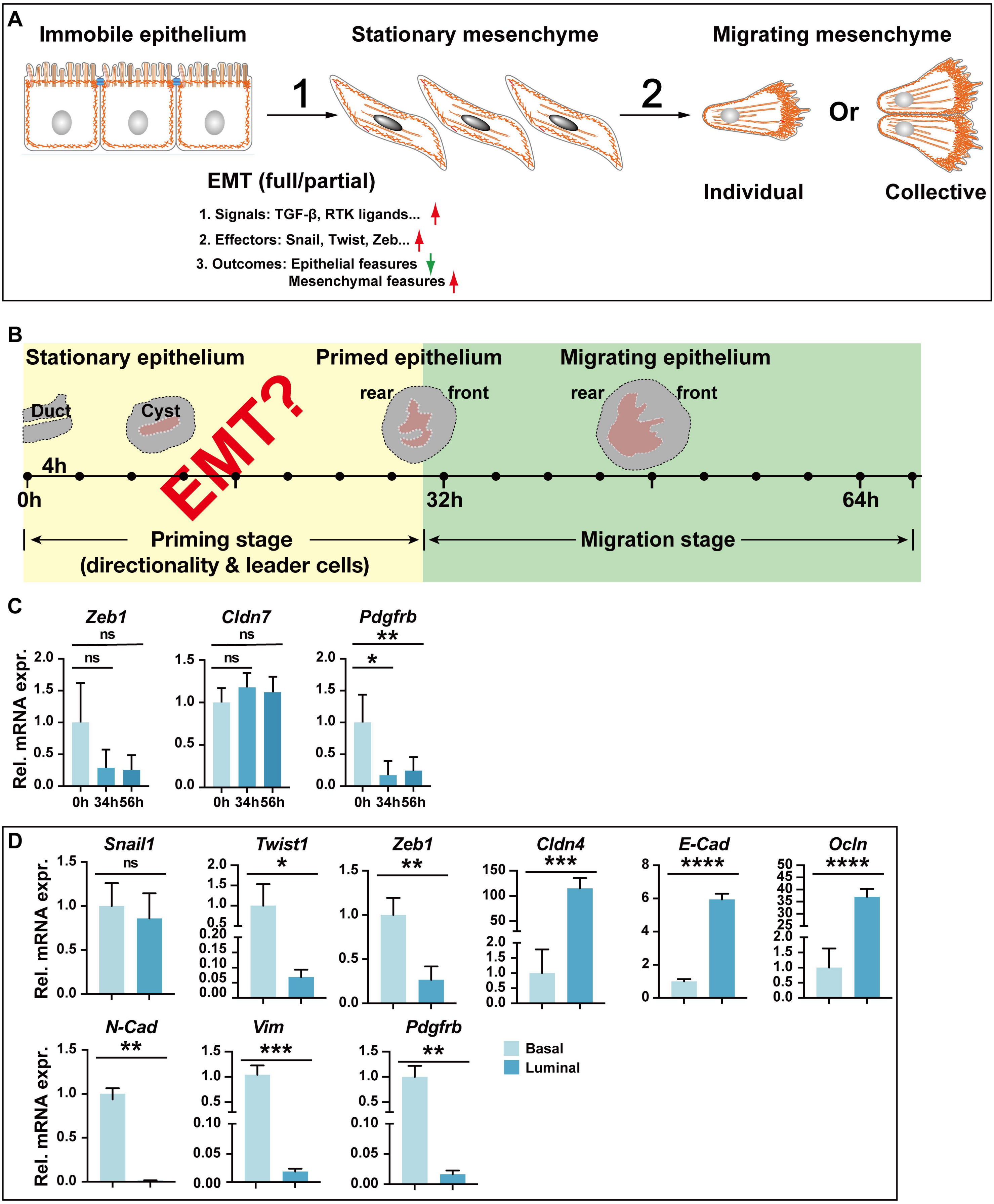

**Sup. Fig. 2. mRNA Expression of Cell-Cell Adhesion Components Does Not Change as Predicted by Classical EMT Models.**

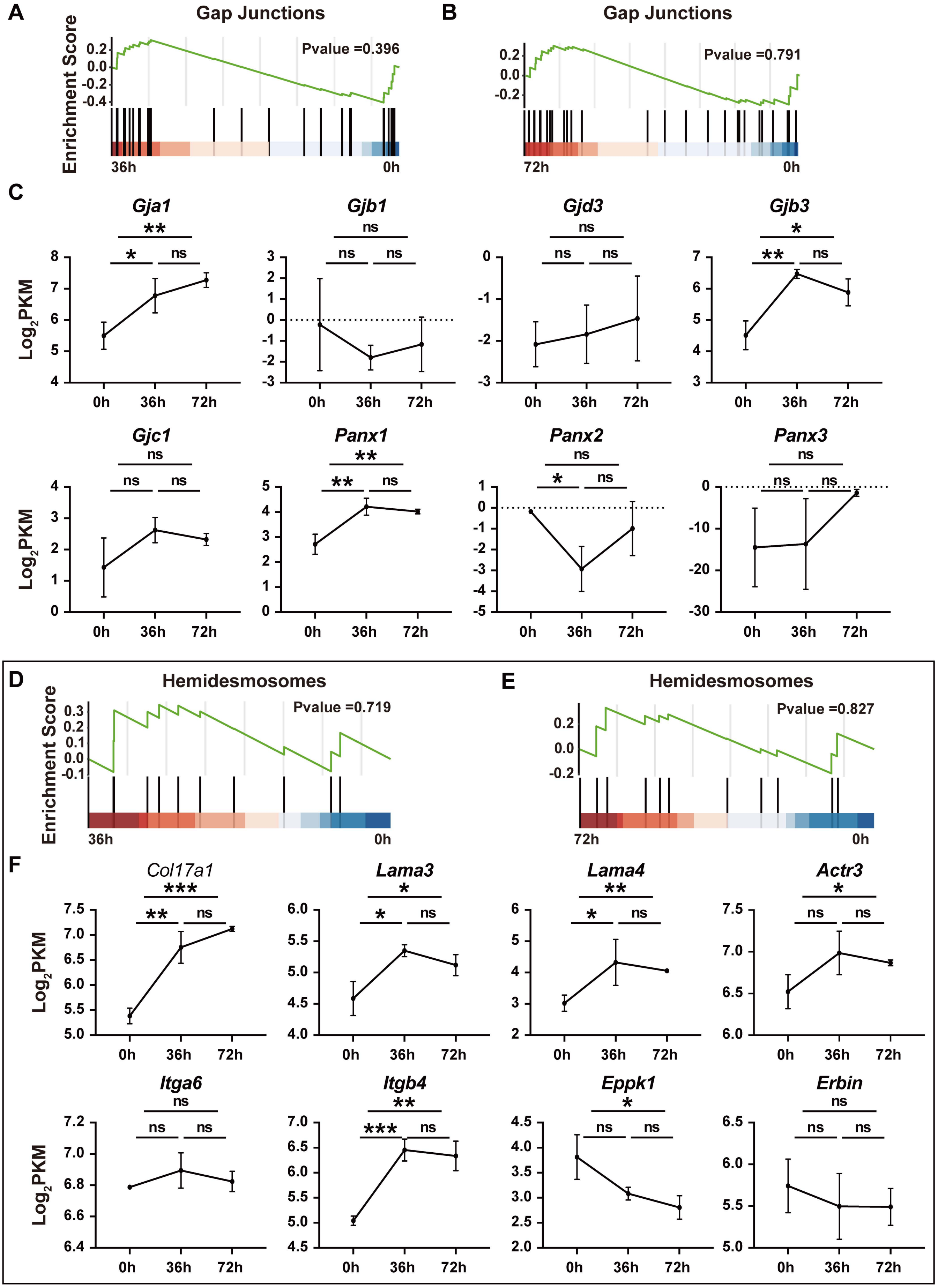

**Sup. Fig. 3. mRNA expression of genes in TJs and desmosomes is reduced and increased, respectively.**

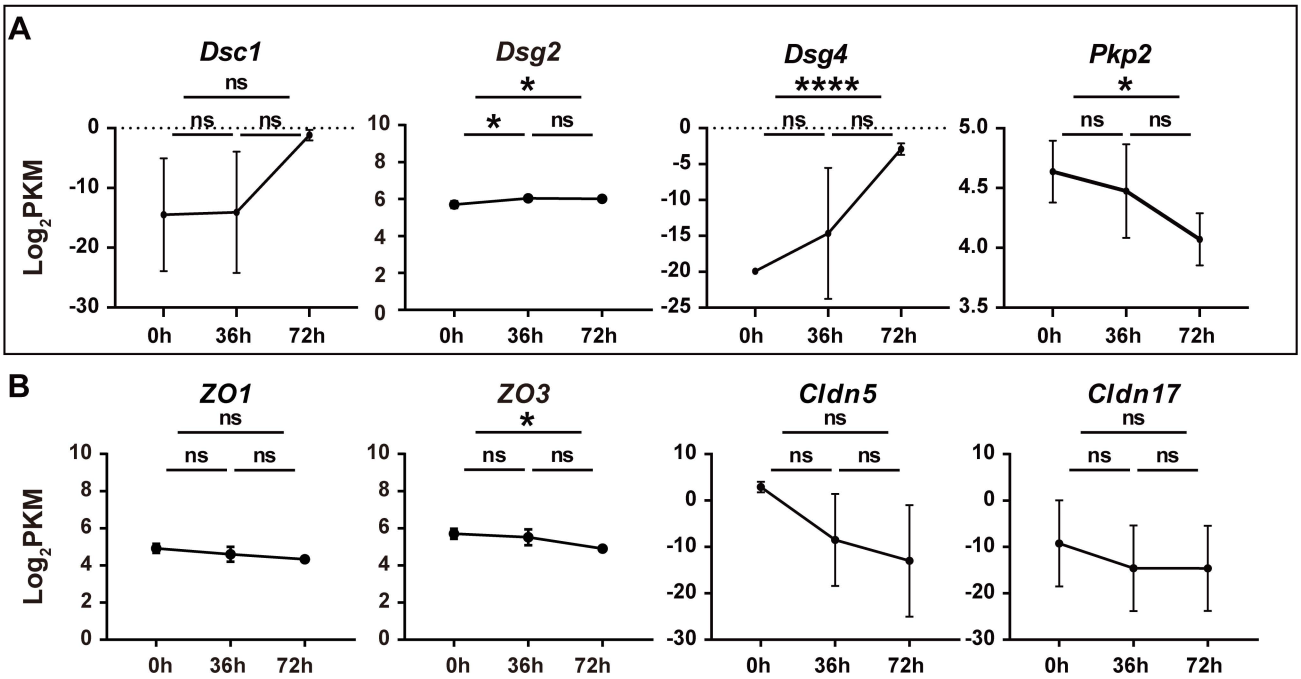

**Sup. Fig. 4. Tight Junctions Are a Barrier to, But Cell-Cell Adhesion is Required for, Epithelial Migration.**

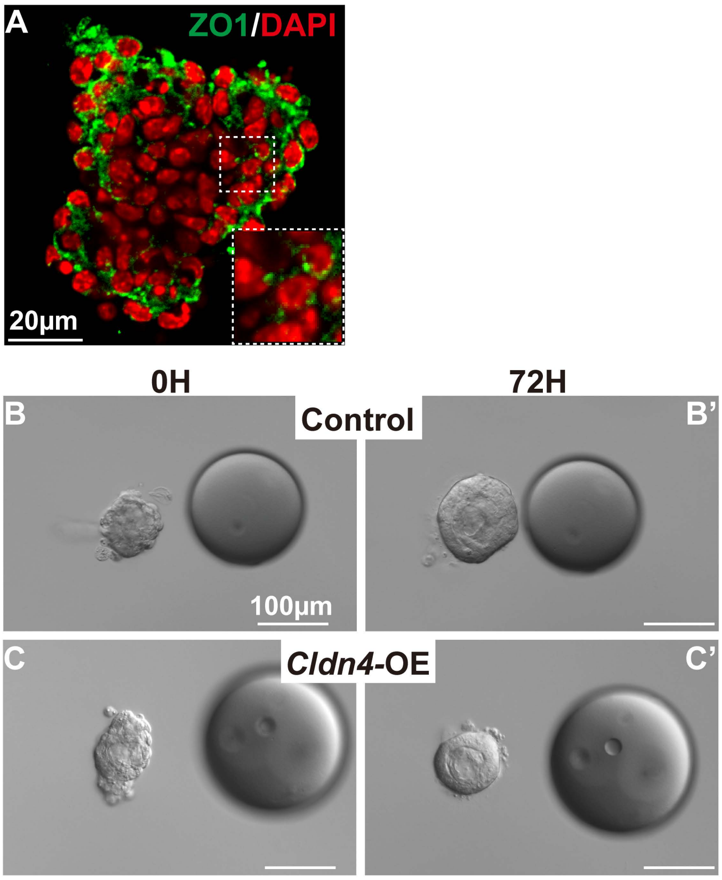

**Sup. Fig. 5. *Snail1* promotes cell-cell adhesion and multiple aspects of directional migration.**

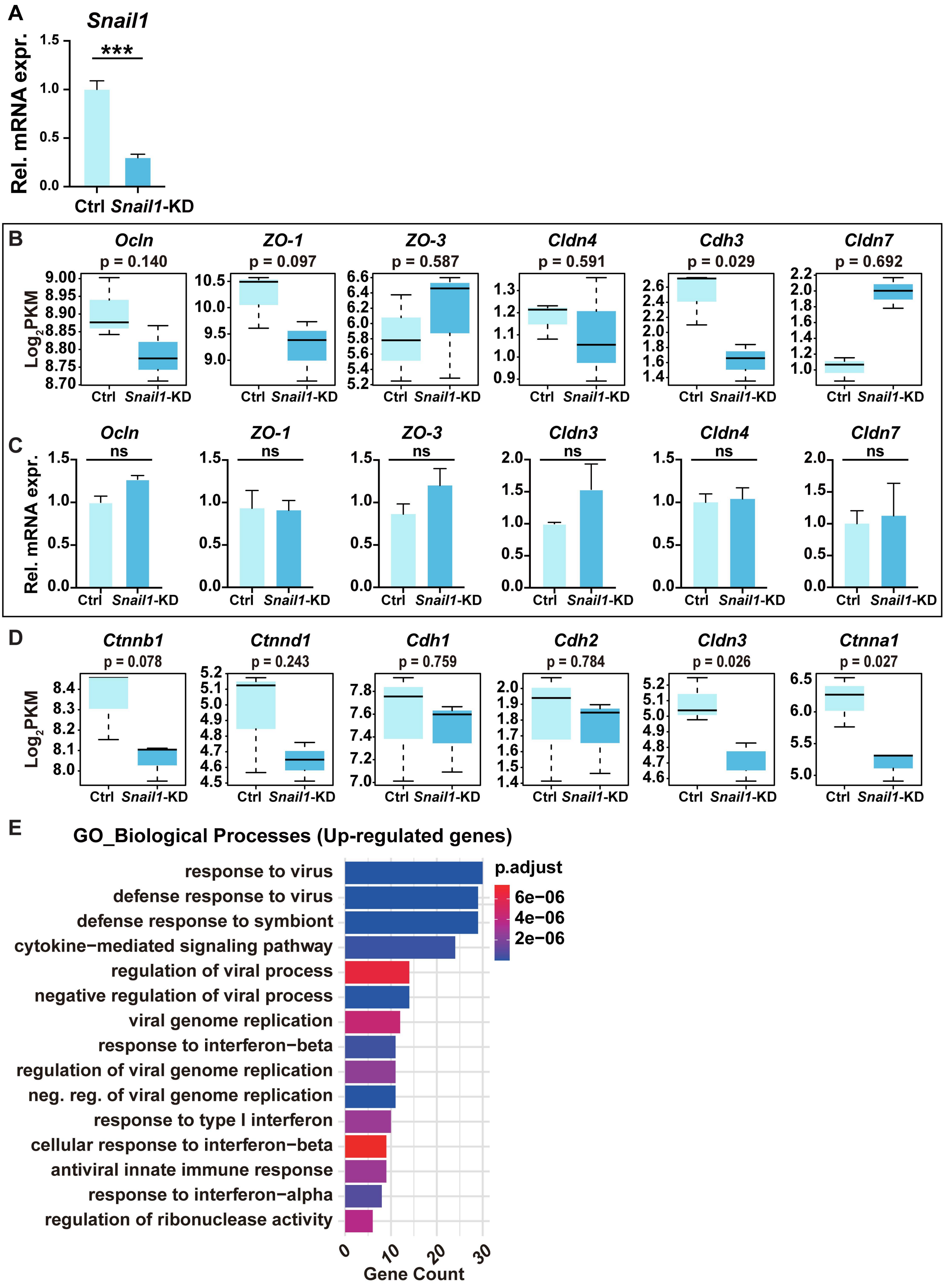

**Sup. Fig.6. *Ep5* regulates moving direction during epithelial migration.**

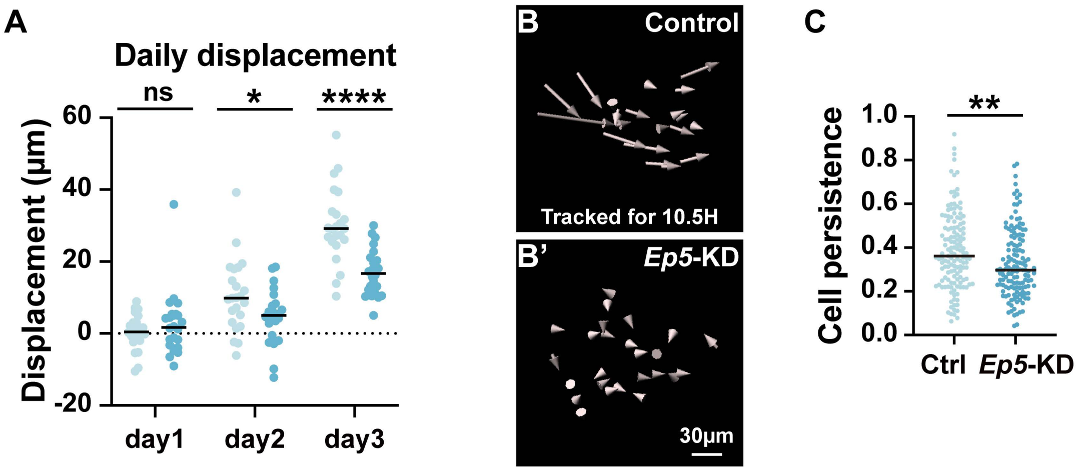

**Supplementary Figure 1: Mammary epithelium does not show typical EMT signatures during collective migration.**

(**A**) Diagram of two stages of epithelial migration based on EMT models: first, static epithelium having apical-basal polarity receives EMT signals or triggers such as TGF-β or RTK ligands, which then activates EMT transcription factors, including *Snail*, *Twist*, and *Zeb*, leading to a decrease of epithelial features like loss of cell-cell/cell-matrix adhesions, and gain of mesenchymal features including emergence of individual or loosely connected cells enriched in mesenchymal marker expression, including *N-Cadherin*, *Vimentin* expression. Note, EMT transition could be complete, or hybrid where an intermediate state expressing both epithelial and mesenchymal markers; second, fully or incompletely transitioned mesenchymal cells migrate individually or, as emerging evidence suggests, collectively.

(**B**) Diagram of two stages of epithelial migration based on recent studies [15]; first, FGF10 gradient promotes preferential cell proliferation in the front of stationary organoid epithelium, and sets up the ‘‘front-rear’’ polarity. As a result, the organoid becomes primed with the front being stratified and converting apicobasal polarity to front-rear polarity, i.e. gaining directionality. Second, organoid epithelium becomes migratory and moves toward FGF10 signal.

(**C**) Relative mRNA expression of select EMT markers as detected by qPCR at different stages of mammary epithelial migration.

(**D**) Relative mRNA expression of a panel of EMT markers as detected by qPCR in primary basal and luminal cells.

Data are mean ± SD. Statistical analysis was performed using unpaired Student’s t test. *$p < 0.05$; **$p < 0.01$; n.s., not significant. Abbreviations: Rel, relative; expr, expression.

**Supplementary Figure 2: mRNA expression of genes in TJs and desmosomes is reduced and increased, respectively.**

(**A, B**) GSEA of gap junction genes between organoids at 36 hours and 0 hour (**A**) and 72 hours and 0 hour (**B**). Note the P values were not significant.

(**C**) Relative mRNA expression of gap genes at the indicated stages of mammary organoid migration.

(**D, E**) GSEA of hemidesmosomes genes between organoids at 36 hours and 0 hours (**D**) and 72 hours and 0 hours (**E**). Note the P values were not significant.

(**F**) Relative mRNA expression of hemidesmosomes genes at the indicated stages of mammary organoid migration.

Data are mean ± SD. Statistical analysis was performed using unpaired Student's t test. $*p < 0.05$; $**p < 0.01$; $***p < 0.001$; $****p < 0.0001$; n.s., not significant.

**Supplementary Figure 3: mRNA expression of genes in TJs and desmosomes is reduced and increased, respectively.**

(**A, B**) Relative mRNA expression of select genes of desmosomes (**A**) and tight junctions (**B**) at the indicated stages of mammary organoid migration.

Data are mean ± SD. Statistical analysis was performed using unpaired Student's t test. $*p < 0.05$; $****p < 0.0001$; n.s., not significant.

**Supplementary Figure 4: Tight junctions are a barrier, but cell-cell adhesion is required for epithelial migration.**

(**A**) Immunofluorescence of ZO1 on MEC aggregates from primary basal and luminal cells co-stained with the nuclear dye DAPI. Note the aggregates lack TJs, as indicated by the lack of a lumen and characteristic belt-like ZO1 signals associated with TJs (**A'**). Scale bars, 20 µm.

(**B**, **C**) DIC images of the time course of mammary epithelial cell aggregates expressing either a control vector (**B**, **B'**) or *Cldn4* (**C**, **C'**) at the time-points indicated. Scale bars, 100 µm.

**Supplementary Figure 5: *Snail1* promotes cell-cell adhesion and multiple aspects of directional migration.**

(**A**) Relative *Snail1* mRNA expression in HC11 cells transfected by the control or Snail1-KD lentiviral construct.

(**B**, **C**) mRNA expression of a panel of TJs components based on the *Snail1*-KD bulk RNA-sequencing data (**B**) and qPCR validation (**C**).

(**D**) mRNA expression of a panel of adherence junctions components based on the *Snail1*-KD bulk RNA-sequencing data.

(**E**) GO analyses based on top 100 most highly down-regulated genes in *Snail1*-KD HC11 aggregates when compared with the control aggregates. Gene ontology analysis of the differentially ($p < 0.05$) expressed genes to determine the biological processes with which these genes might be involved.

Data are mean ± SD. Statistical analysis was performed using unpaired Student's t test. ***, $p < 0.001$; N.S., not significant. Abbreviations: Rel, relative; expr, expression.

**Supplementary Figure 6: Cdc42ep5 regulates moving direction during epithelial migration.**

(**A**) Quantification of daily displacement of epithelial migration of the control (n = 23) or *Ep5*-KD (n = 24) HC11 aggregates toward beads pre-soaked in FGF10.

(**B**-**C**) Persistence of individual cells from the control (n of aggregates = 6, n of tracks =139) and *Ep5*-KD (n of aggregates = 5, n of tracks =139) aggregates.

Data are mean ± SD. Statistical analysis was performed using unpaired Student's t test. *$p < 0.05$; **$p < 0.01$; ****$p < 0.0001$; N.S., not significant.
